# Development of an eco-friendly RNAi yeast attractive targeted sugar bait that silences the *Shaker* gene in spotted-wing drosophila, *Drosophila suzukii*

**DOI:** 10.1101/2025.03.26.645612

**Authors:** Keshava Mysore, Jackson Graham, Teresia M. Njoroge, Akilah T. M. Stewart, Saisuhas Nelaturi, Molly Duman-Scheel

**Affiliations:** Department of Medical and Molecular Genetics, Indiana University School of Medicine, Raclin-Carmichael Hall, South Bend, IN 46617 USA; Eck Institute for Global Health, University of Notre Dame, Notre Dame, IN, 46556, USA; Department of Biological Sciences, University of Notre Dame, Notre Dame, IN, 46556, USA; Department of Chemistry, University of Notre Dame, Notre Dame, IN, 46556, USA

**Author notes:** Corresponding author: Molly Duman-Scheel, Indiana University School of Medicine, Raclin-Carmichael Hall, South Bend, IN 46617.

**Keywords:** fruit, crop, integrated pest management, insecticide, SWD

## Abstract

**Background:** *Drosophila suzukii,* or spotted-wing drosophila (SWD), (Diptera: Drosophilidae), are invasive vinegar flies of East Asian origin that have wreaked havoc on the small fruit and berry industry. In locations where SWD are well established, weekly chemical insecticide applications are necessary, resulting in increased economic costs, unwanted environmental impacts ensuing from loss of non-targeted organisms, and the eventual emergence of populations that are resistant to these insecticides. It is therefore critical that new classes of biorational pesticides and cost-effective technologies for controlling SWD are identified.

**Results:** Here, we used the attractive properties of Saccharomyces cerevisiae, baker’s yeast, which was designed to express an RNA interference (RNAi) pesticide that specifically targets the SWD *Shaker (Sh)* gene, to lure and kill flies that feed on the yeast, which was delivered in a feeder as a component of an attractive targeted sugar bait (ATSB). The yeast, which was heat killed prior to preparation of the ATSB, silenced the *Sh* gene, resulting in severe neural defects and 96±9% fly mortality in laboratory trials. The RNAi yeast was successfully fed to the flies in an easily assembled soda bottle feeder that continuously rewetted the yeast with soda, which lured and killed the flies in simulated field trials. Despite this toxicity observed in SWD, consumption of the yeast had no impact on the survival of other dipteran insects.

**Conclusion:** This promising ATSB technology, which was prepared with a new class of RNAi yeast insecticides, could one day be an effective component in integrated SWD control programs.

## 1. Introduction

*Drosophila suzukii,* or SWD, are invasive vinegar flies of East Asian origin. SWD, which lay eggs in a variety of wild and cultivated plants, have decimated the small fruit industry worldwide, including Europe, the Americas and Africa^1–4^. SWD, which complete multiple generations in a year, impact most berry crops, cherries, grapes and other tree fruits^5^, generating upward of 80% crop loss, and an estimated $700 million economic loss for producers annually when they are not adequately controlled (https://shorturl.at/VLhNF). SWD females penetrate the skin of fresh fruit prior to harvest, laying eggs under the skin, and compromising the fruit integrity, resulting in dimpling, wrinkling, and browning of the larval-infested fruit as well as potential sites for infestation by other insects and infection sites for microbial pathogens^2^. In regions where SWD are well established, weekly insecticide applications are necessary^6,7^, generating increased economic costs, as well as unwanted environmental impacts resulting from loss of non-targeted organisms, such as bees, butterflies, and other pollinators. With increased use of insecticides, SWD resistance to organophosphates and pyrethroids is a major threat^8,9^. Moreover, once eggs have been deposited, insecticidal sprays will no longer protect the fruit, which has been infested with maggots^2^. It is therefore important to find means of targeting adult female flies to prevent egg-laying behavior. Here, we evaluate a new environmentally-friendly species-specific insecticide that can be deployed as an attractive targeted sugar bait (ATSB) to combat SWD infestations.

ATSBs utilize the natural sugar feeding behavior of insects such as mosquitoes and flies, many of which require plant sugar for survival and reproduction. ATSBs, which have been developed for the control of vector mosquitoes, capitalize on this natural sugar feeding behavior to attract mosquitoes that feed on a sugar source that has been laced with a poison^10^. Although ATSBs have not yet been assessed broadly in SWD, trials have demonstrated that ATSBs can significantly reduce sand fly^11^ and mosquito^12^ populations. Sugar baits facilitate targeted delivery of a variety of pesticides^10^, resulting in an overall reduction in pesticide applications, but resistance is nevertheless still of concern. Furthermore, first-generation mosquito ATSBs have centered on the use of non-specific chemical insecticides that can harm non-target insects^13^. To address these concerns, we recently identified a new class of RNA interference (RNAi) pesticides that result in significant mortality in adult mosquitoes of a variety of species when delivered as ATSBs in lab^14,15^ and semi-field^16^ trials. Use of RNAi insecticides, which are species-specific, enhances existing ATSB technologies and may represent a new generation of insecticides to combat insecticide resistance.

In the RNAi pathway, a conserved mechanism that helps protect organisms from viral infections, small interfering RNAs (siRNAs) silence (turn off) genes that are complementary in sequence^17^. This complementarity facilitates the design of custom insecticides^18^. We have developed multiple RNAi pesticides that target neural genes in various types of mosquitoes^14,15^. These mosquito-specific insecticides were designed to target gene sequences conserved in mosquitoes, but not in any other organisms, including humans. Interfering RNA can be expressed in *S. cerevisiae*, a model organism that is genetically tractable, inexpensive to culture at scale, and which can facilitate cost-effective RNA production during yeast cultivation^19^. The yeast, which can be heat killed prior to use, is highly attractive to adult mosquitoes, which can consume it through ATSBs delivered in bait station sachets^16^. Yeast RNAi systems promote high levels of gene silencing in adult mosquitoes, and when used to target genes that are required for mosquito survival, the yeast can generate high levels of mortality upon consumption^14,15^. Moreover, the yeast formulations developed for mosquito control were shown to maintain activity for several months^20^, suggesting that a single deployment of yeast ATSBs (rather than weekly treatments) could be sufficient to control SWD throughout a single growing season in the north central United States. Here, we aimed to extend this RNAi yeast ATSB research to develop a new SWD control intervention.

Several of the genes that were successfully targeted by mosquito RNAi pesticides are also present in SWD flies. One example is the *Sh* gene, which functions in the mosquito brain^21^. *S. cerevisiae* (baker’s yeast) was engineered to express interfering RNA corresponding to mosquito *Sh* genes. *Sh* genes encode an evolutionarily conserved subunit of a voltage-gated potassium channel, a regulator of neural activity in both invertebrate and vertebrate organisms^22,23^. Consumption of the yeast by mosquitoes silenced *Sh*, resulting in the death of adult females that consumed yeast delivered through a sugar bait. Mortality correlated with defects in the mosquito brain. The insecticidal activities of the yeast were subsequently confirmed in trials conducted on *Aedes albopictus*, *Anopheles gambiae*, and *Culex quinquefasciatus* mosquitoes, yet the yeast was not toxic to non-target arthropods that lacked the *Sh* target sequence^16,21^. These studies indicate that yeast RNAi pesticides targeting *Sh* could be further developed as broad-based mosquito insecticides for utilization in integrated biorational mosquito control programs. These findings indicated that the species-specificity of ATSBs, a new paradigm for vector control^10^, could be enhanced through the use of RNAi yeast-based pesticides.

Although the mosquito yeast RNAi pesticides developed to date do not have a target site in *D. suzukii,* a copy of the *Sh* gene is found in the SWD genome^24^. We hypothesized that modification of the shRNA sequence such that it matches the SWD version of the *Sh* gene, but not that of other organisms, would allow us to specifically kill SWD. Likewise, recent work^25^ demonstrated that baker’s yeast is a potent attractant for SWD in the field, where it performs superiorly to other attractants. Here, we evaluate RNAi yeast targeting the SWD *Sh* gene by incorporating it into an ATSB that is fed to SWD.

## 2. Materials and Methods

### 2.1 Insect rearing

An SWD strain prepared from a local Michigan collection was obtained from Juliana Wilson (Michigan State University) and reared in a 26° C,lrJ∼80% relative humidity, with a 12 h light/12 h dark cycle with 1 h crepuscular periods. Flies were reared in bottles containing Nutri-Fly® BF, (Genesee Scientific, El Cajon, CA) media.

### 2.2 Preparation and culturing of Sh.706 yeast

An RNAi yeast strain corresponding to the D. suzukii potassium voltage-gated channel protein-encoding gene Sh (NCBI Gene ID: 108016735) target site CGTTGAGCGGTGAAAACCTATCCAA was generated as previously described^26^ using the S. cerevisiae CEN.PK yeast strain (genotype MATa/α ura3-52/ura3-52 trp1-289/trp1-289 leu2-3_112/leu2-3_112 his3 Δ1/his3 Δ1 MAL2-8C/MAL2-8C SUC2/SUC2)^27^. The yeast was transformed with a pRS426GPD plasmid^28^ bearing the Sh shRNA expression cassette, which was prepared as described^26^. Transformants were selected through growth on media lacking uracil. The yeast, hereafter referred to as Sh.706, was cultured and heat killed upon harvesting as described^26^, then freeze dried with 0.025% benzoic acid (as a preservative).

### 2.3 Initial ATSB screening assays

3-4 day old flies were starved for 4-5 hours by transferring them into an empty bottle. Control and treatment yeast were prepared. At the end of the starvation period, the bottle of flies was placed on ice for 15-20 min. 100 µl of 10% sucrose solution with 4.5 % red food coloring was mixed with 40 mg of yeast using a sterile toothpick. Four ∼25 µl drops of the sugar+yeast mixture were placed on a sterile 100 mm x 15 mm petri dish. 25 flies were placed on the petri dish, which was then covered with its lid. Flies were then placed at room temperature (∼21±1°C) for overnight feeding. Feeding was confirmed by the presence of red food dye in the abdomens of the flies. The next morning, flies from each treatment were again placed on ice and then transferred to fresh food vials, in which they were monitored for six days. Behavioral phenotypes and survival were assessed. The data were tabulated, analyzed using the Student’s t-test, and graphed in Microsoft Excel 365 software. The survival curves were generated using SPSS 25 (IBM, Armonk, NY USA) software by applying the Kaplan-Meier method.

### 2.4 ATSB feeding experiments

Modified MUDUODUO automatic bird drinker cups (Amazon, Seattle Washington) were used to feed yeast to the flies. A small piece of clean dehumidifier filter (Honeywell Home, Charlotte, NC) without the metal layer was inserted along the channel between the feeding area and the reservoir. The channel and reservoir were wrapped with parafilm and covered to minimize leakage. At the feeder end, a platform wick was created using the same filter cut into a 3.3 cm x 7.0 cm rectangle and folded into a circle. Food grade nylon membranes (Amazon, Seattle Washington) of 5 µm (bottom) and 90 µm (top) were layered on top. A paste was prepared by mixing 40-200 mg of either control or treatment yeast with 200 µL of degassed/flat Coca-Cola and placed between the membrane layers. A soda bottle (12 Fl OZ) with 110 mL of Coca-Cola and 20 ml/L Tegosept anti-mold agent (Thermo Fisher Scientific, Waltham, Massachusetts) was inverted to act as a continuous supply of soda (Supplemental Fig. 1). This feeder along with 50 3-4 day old sugar-starved flies were placed in insect cages. The experiment was set up in the insectary, where it was monitored for six days. Mortality and any other phenotypic changes were noted daily. The data were tabulated and graphed in Microsoft Excel 365 software. One-way ANOVA statistics were performed and survival curves generated using SPSS 25 (IBM, Armonk, NY, USA) software. Kaplan-Meier method was utilized for the survival curves.

### 2.5 Confirmation of *Sh* gene silencing

Silencing of the SWD *Sh* gene was examined through qRT-PCR. Pools of five whole adult flies fed with Sh.706 as described above were collected 16 hrs after yeast-ATSB feedings (control or treatment). TRIzol (Invitrogen, Carlsbad, CA) was used to extract total RNA as described in the manufacturer’s instructions; the RNA was then treated with DNase using the DNA-free DNA Removal kit (Invitrogen, Thermo Fisher Scientific, Waltham, MA) according to the manufacturer’s instructions. The High Capacity RNA to cDNA Kit (Applied Biosystems, Foster City, CA) was used for production of cDNA, which was amplified in an Applied Biosystems Step One Plus Real-Time PCR System using the Power SYBR Green PCR Master Mix (Applied Biosystems, Foster City, CA) and the following primers: Sh1 for: 5’ ATCGAGGAAGACGAGGTGCC 3’ and Sh1 rev: 5’ TCCTCCTCTTCGGCAACGAC 3’. Amplification of *alpha tubulin* was performed with the primers 511 AGGATGCGGCGAATAACT 311 (forward) and 511 CGGTGGATAGTCGCTCAA 311 (reverse)^29^ and used for data standardization. PCR reactions were performed in six replicate wells in each of three separate biological replicate trials. Results from these assays were quantified through standardization of reactions to levels of *alpha tubulin* using the ΔΔCt method as described^30^. Data were statistically analyzed using Microsoft Excel 365 software with Student’s t-test.

### 2.6 Immunohistochemistry studies

Immunohistochemical staining of adult fly brains was performed after Sh.706 treatments using the following reagents as described^31,32^: mAb nc82 anti-Bruchpilot^33,32^ (DSHB, Iowa Hybridoma Product nc82 deposited by E. Buchner), anti-HRP (Jackson ImmunoResearch Labs, West Grove, PA), and TO-PRO-3 iodide (Molecular Probes, Eugene, OR). The staining assays were performed in triplicate, and stained tissues were mounted and imaged using a Zeiss 710 confocal microscope. Confocal images were analyzed with FIJI ImageJ^34^ and Adobe Photoshop 2025 software; mean gray values (average signal intensity over the selected area) were calculated as previously described^35^ and compared using one-way ANOVA and Bonferroni post-hoc test in SPSS 25 (IBM, Armonk, NY, USA) software. The data were graphed in Microsoft Excel 365 software.

### 2.7 Dose-response curves

Dose-response assays were conducted as described in mosquitoes^36^ through the generation of different concentrations of ATSB yeast using varying amounts of control and treatment RNAi yeast. The experiments were carried out with 25 individuals/treatment in a petri dish as described above. The final data were tabulated, and probit analyses were performed with SPSS 25 (IBM, Armonk, NY, USA) software. The data were graphed in Microsoft Excel 365 software.

### 2.8 Non-target assays

The impacts of Sh.706 yeast feedings on non-target dipterans were assessed in *Drosophila melanogaster, Aedes aegypti, Anopheles stephensi,* and *Culex quinquefasciatus*. Insects were reared, and yeast sugar feeding assays were performed as described^14^.

## 3. RESULTS

### 3.1 Adult fly consumption of yeast expressing shRNA targeting SWD *Sh* results in insect mortality

RNAi yeast expressing shRNA corresponding to the SWD *Sh* gene was prepared and characterized. Although the Sh.706 yeast corresponds to a target sequence present in the *D. suzukii* and *D. melanogaster Sh* genes, NCBI blast searches did not identify other organisms with an identical 25 bp target site. Following culturing, heat-inactivation, and drying of the yeast, it was mixed with sucrose and offered to SWD. Although SWD fed with sucrose containing a control yeast strain^36^ that expresses shRNA with no known target in SWD survived, significant mortality (96±9%, P<0.001 vs. control) was observed within six days after SWD consumed the Sh.706-sucrose bait (Fig. 1A,C). Dose-response assays for Sh.706 generated the curve shown in Fig. 1D, which revealed an LD_50_ of 189 µg yeast/µL of 10% sucrose.

**Figure 1.**
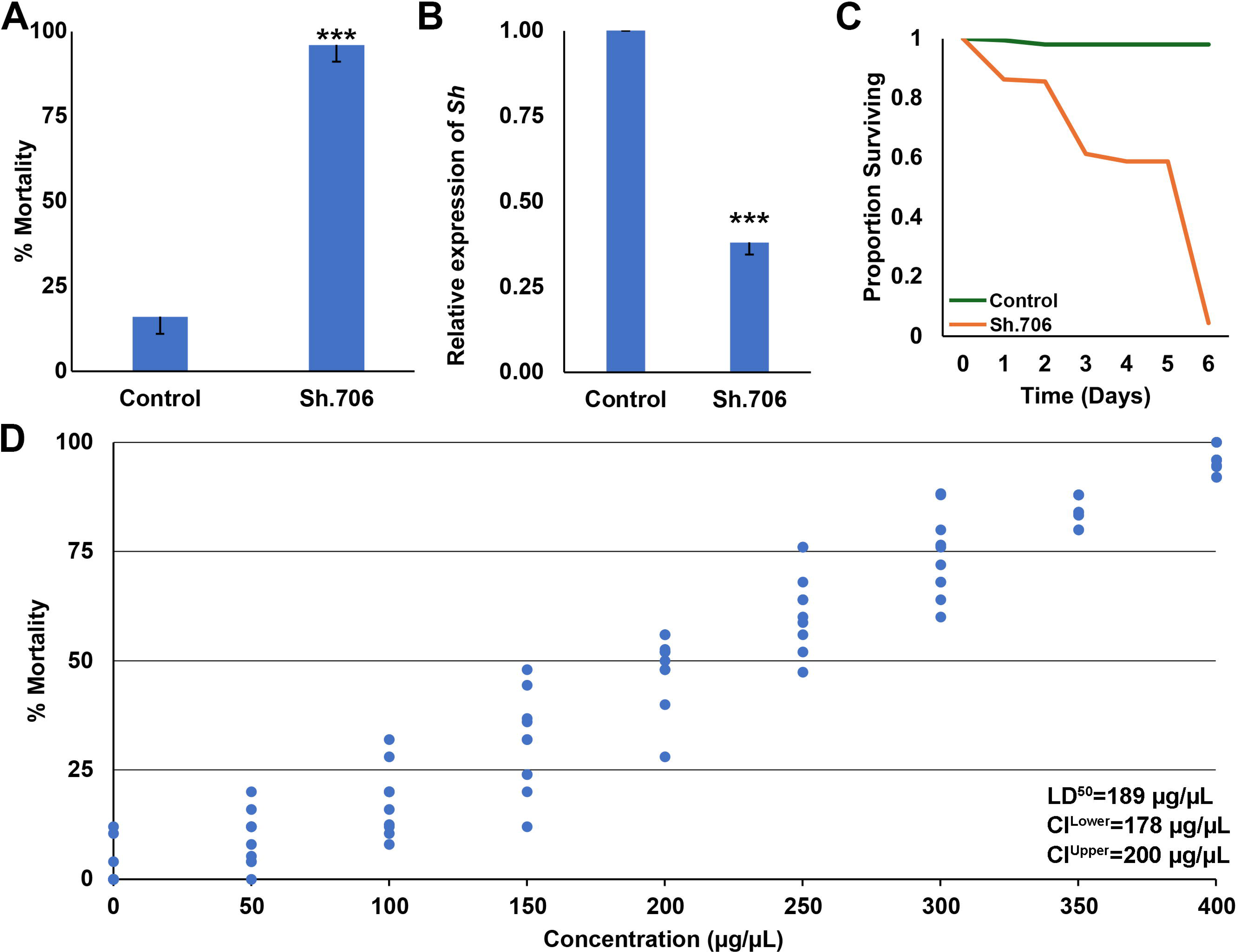
Insecticidal activity of Sh.706 yeast. (A) Laboratory experiments showed that yeast expressing Sh.706 shRNA caused significant mortality in SWD flies compared to those fed with yeast expressing control shRNA with no target specificity (***P < 0.001, Student’s t-test; error bars represent the standard error of the mean, SEM). (B) qRT-PCR confirmed that the SWD *Shaker* gene was silenced in flies that consumed Sh.706 yeast compared to those fed with control yeast (***P < 0.001, Student’s t-test; error bars represent SEM); a survival curve is shown in (C). (D) Dose-dependent mortality was observed in *D. suzukii*, with an LD_50_ of 189 µg/µL. Data were compiled from ten replicate trials per condition, each containing 25 adults.

### 3.2 Silencing of *Sh* expression results in neural and behavioral phenotypes in adult flies

It was hypothesized that neural defects would be observed in SWD flies that had consumed the RNAi yeast ATSB. 60% silencing of *Sh* expression, which was verified through qRTPCR (Fig. 1B), correlated with neural (Fig. 2) and behavioral defects. *Sh* silencing correlated with 58% reduction in levels of nc82, which labels Bruchpilot, a marker of active synapses^32^, in the adult brain (Fig. 2A1 vs B1 and C, green); in addition, the anti-HRP labeled brains showed ∼80% reduction in expression when compared with control brains (Fig. 2A2 vs B2 and C, red). However, no significant differences in TO-PRO nuclear staining were observed in treated and control fly brains (P>0.05, Fig. 2A3 vs B3 and C, blue). Shaking was observed in the legs of treated flies, which had locomotor defects and were unable to fly.

**Figure 2.**
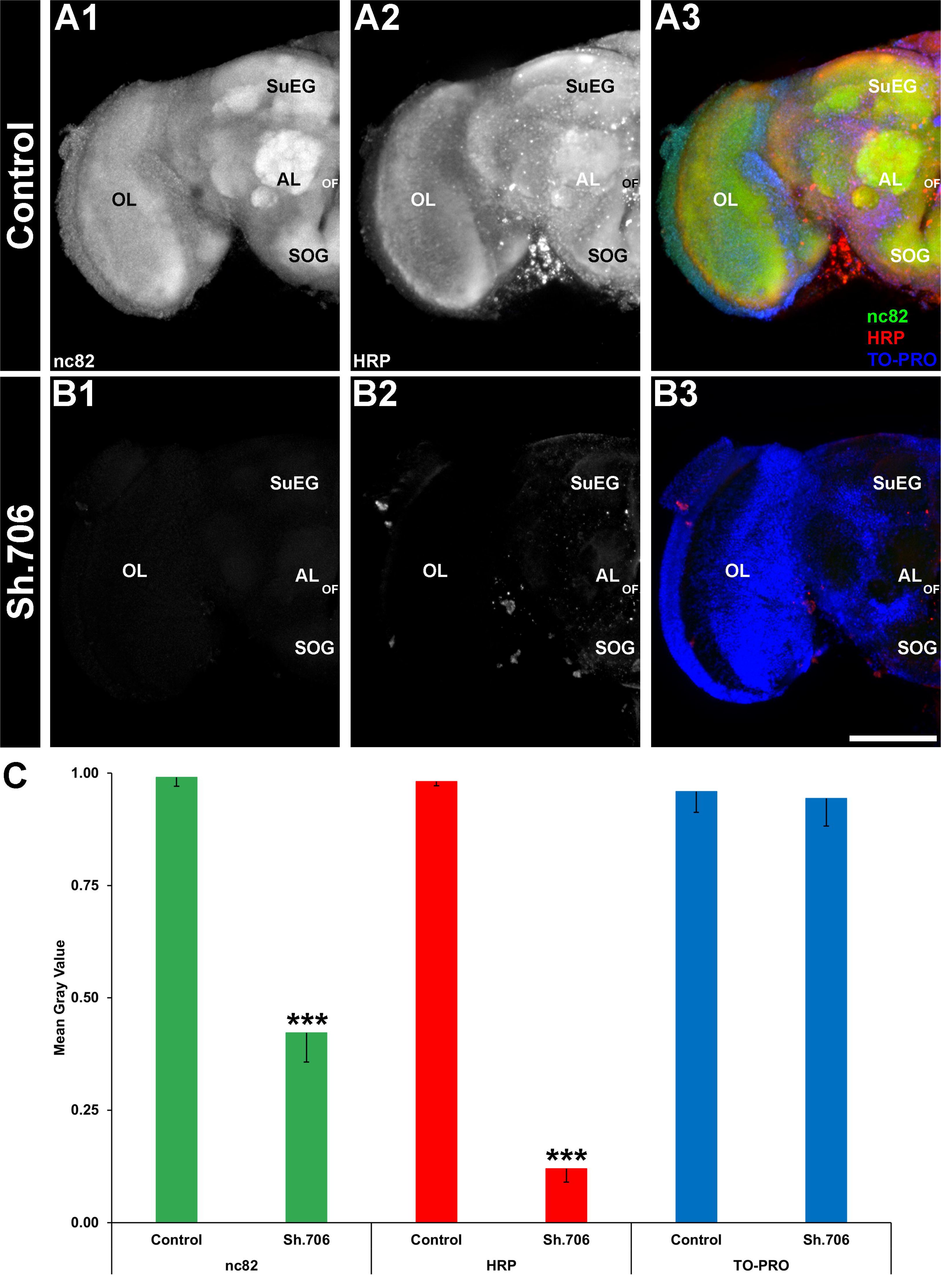
Neural defects in *D. suzukii* adults fed with Sh.706 yeast. Adult brains were labeled with three antibodies: mAbnc82 (active synapse marker; white in (A1, B1); green in (A3)), anti-HRP antibody (neural marker; white in (A1, B2); red in (A3)), and TO-PRO (nuclear marker; blue in (A3, B3)). The levels of nc82, a protein involved in neural development, were reduced by ∼50% in the synaptic neuropil of adults fed with Sh.706 yeast compared to control yeast-treated adults (A1 vs B1, C-green bars). The general neural marker HRP also showed reduction in expression (∼80% reduction) compared to control-fed individuals (A2 vs B2, C-red bars). Data are presented as average mean gray values, with error bars denoting the standard error of the mean. ***Indicates a statistically significant difference from the control group (P < 0.001). Representative larval brains are oriented dorsal upward. AL, larval antennal lobe; OL, optic lobe; SOG, sub-esophageal ganglion; SuEG, supraesophageal ganglion. A combined total of 57 control and 63 Sh.706-treated individual brains from three biological replicate experiments were analyzed. Scale Bar = 100 μm.

### 3.3 Sh.706 yeast consumption does not impact non-target dipterans

Despite the significant neural and behavioral defects observed in SWD, it was hypothesized, on the basis of the lack of identical target sites in other organisms, that consumption of Sh.706 yeast would not impact survival of non-target dipterans. No significant death (P>0.05) was detected in the survival of adult *A. aegypti, Anopheles stephensi,* and *Culex quinquefasciatus* adult mosquitoes or *D. melanogaster* (which lack systemic RNAi) that consumed Sh.709 yeast or control yeast (Fig. 3), all of which survived.

**Figure 3.**
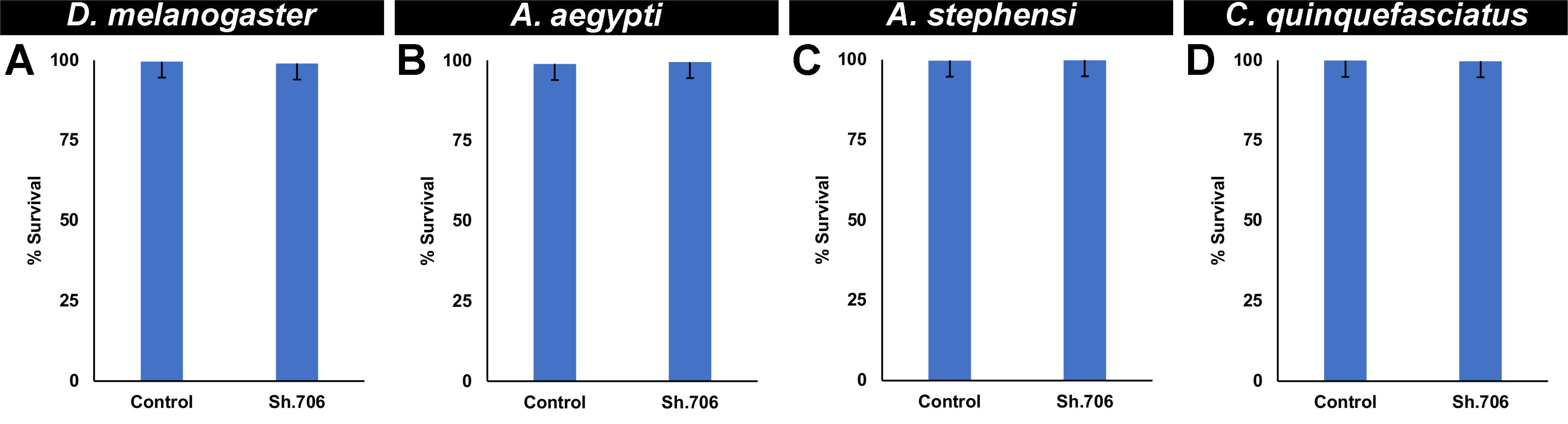
Survival of non-target arthropods consuming Sh.706 yeast. Consumption of Sh.706 yeast by adults of the fruit fly *D. melanogaster* (A) and mosquitoes *A. aegypti* (B), *A. stephensi* (C), and *C. quinquefasciatus* (D) did not affect non-target insect survival. A total of 150 adults per treatment were assessed in two biological replicate experiments for all species. Graphs display mean survival percentages with standard deviation as error bars. Student’s t-test was used to compare means of control-fed vs treatment-fed individuals.

### 3.4 Development of an RNAi yeast ATSB bait station

Soda has previously been shown to function as a sugar bait that can attract mosquitoes^37^. In an effort to create a yeast delivery system that could be easily constructed and evaluated in the field, a Sh.706 yeast-soda feeder system, deemed the yeast everlasting soda (YES) feeder, was prepared (Supplemental Fig. 1, Fig. 4C) and assessed under simulated field conditions (Fig. 4A,B). All flies were observed drinking from the feeders, which were prepared either with soda alone or with soda and Sh.706 or control RNAi yeast. Although flies that drank from the Sh.706 feeder died within six days, flies that consumed the control yeast survived (P<0.001, Fig. 4A,B).

**Figure 4.**
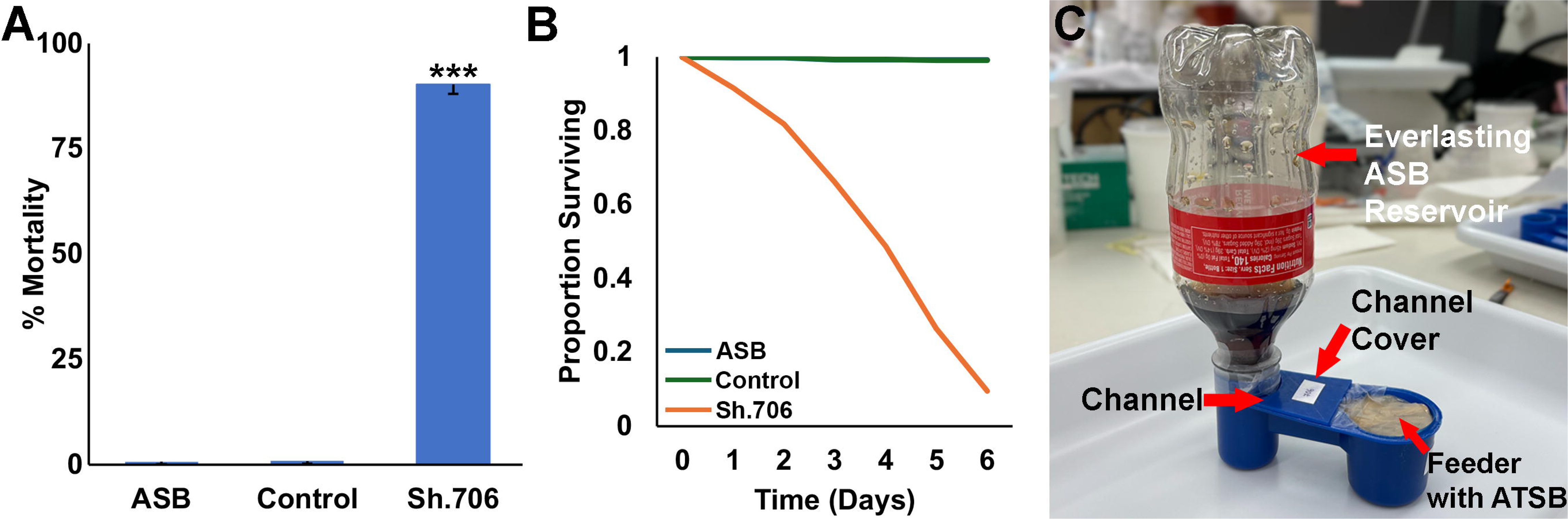
Sh.706 ATSB delivered in a YES feeder induced significant mortality. Sh.706 yeast fed to *D. suzukii* adults as an ATSB prepared with soda and delivered in the YES feeder led to significant mortality when compared with control yeast-fed individuals (A). The YES feeder consists of automatic bird drinker cups modified to deliver yeast ATSB prepared with soda (B). A combined total of 506, 537 and 514 individuals were tested from four biological replicate experiments for ASB, control and Sh.706 treatments, respectively.

## 4. Discussion

Jarausch et al^38^ discussed the great potential, but critical challenges for the development of RNAi control measures for SWD. Among the challenges, they note the need for an effective delivery system for RNAi pesticides. They also discuss the necessity of identifying the proper target genes. In this investigation, an RNAi yeast pesticide that silenced *Sh*, a gene required for SWD survival, was produced at low cost in baker’s yeast, which was heat-inactivated, dried with a preservative, and fed to SWD using attractive sugar baits (Fig. 1). Consumption of the Sh.706 yeast-soda ATSB resulted in lethality which correlated with silencing of *Sh* and defective neural activity in the fly brain, as demonstrated through loss of nc82 and HRP signal that likely results from loss of both neural activity and neural density (Fig. 2). These significant neural defects, accompanied by shaking phenotypes in the flies, correlated with the significant mortality observed in Sh.706-treated flies. It is anticipated that the yeast ATSB, which can be delivered through an inexpensively generated soda bottle feeder system (Fig. 4), could be rapidly deployed and seamlessly integrated with existing SWD strategies, while enhancing these strategies through inclusion of a novel class of SWD-specific pesticides that did not impact non-target dipteran insect survival (Fig. 3). The next step is to evaluate these insecticides in semi-field and field trials to further assess RNAi yeast activity. Given the need to target adult flies prior to egg deposition in fruits^38^, it will be critical to assess the best time and location(s) for deployment of the RNAi ATSB bait stations. It will also be helpful to evaluate how the feeders with soda bait would perform when deployed in outdoor settings where native plants/fruits are present. Furthermore, while the insecticides were not found to be toxic to non-target dipterans, further toxicity testing should be performed with the goal of regulatory authority approvals. If these proposed investigations are successful, RNAi yeast pesticides could be integrated into SWD control programs^39^ with the goal of eradicating this pest in a sustainable and environmentally-conscious manner, improving farmer economic well-being and cost-effective food production.

Scaled production of the yeast would also require the development of robust commercial-ready yeast strains with shRNA expressed from cassettes that have been integrated into the yeast genome, rather than expressed from a plasmid. One method for generating such strains was recently discussed by Brizzee et al.^40^, a study that used Cas-CLOVER and Super PiggyBac engineering in *S. cerevisiae* to integrate multiple copies of an insecticidal shRNA expression cassette into the yeast genome. A thirty-fold increase in shRNA production was observed in the new strain, which reduces the amount of yeast that must be consumed to result in mosquito death and is expected to further reduce the cost of the intervention. Moreover, the Cas-CLOVER strain performed well in pilot scaled fermentations, in which specialized, expensive media were not required, suggesting that scaling yeast production will be both straightforward and inexpensive.

Jarausch et al^38^ also discuss the challenge of addressing biosafety considerations and the importance of avoiding non-target impacts of RNAi pesticides. It should be noted that although other researchers have discussed use of a symbiotic yeast RNAi system that would require the use of live yeast^41^, the yeast pesticides used in this study were heat-inactivated. This is expected to significantly alleviate potential regulatory concerns associated with the release of live RNAi yeast and permits classification of the yeast insecticide as a dead microbial. Moreover, this study is the first to combine the specificity of the RNAi pesticide with the attractiveness of a sugar bait for SWD flies, resulting in a SWD-specific ATSB. A new class of RNAi pesticides could help alleviate insecticide resistance while posing little if any threat to non-target organisms. The use of bait stations also allows the insecticides to be delivered in a targeted manner, which greatly reduces the indiscriminate use of pesticides. Previous baited traps have made use of pheromones to lure SWD^42^. The inclusion of such pheromones could further increase the attractiveness of the ATSBs to SWD. This will be critical in the field, where competitive natural sugar sources are also available. The soda formulations with add mold-inhibitor worked well in lab trials, but it will be critical to assess their efficacy in the field. In the event that these lures do not perform well, other lures could also be assessed. For example, a quinary blend that is both efficient and selective for SWD was identified^43^ and could be assessed in conjunction with RNAi yeast. In the future, SWD repellents could potentially be combined with the ATSB technology to develop push-pull strategies^44^. Finally, the development of a spray formulation, which could be applied to competing non-crop sugar sources, could also prove useful. *Bacillus thuringiensis israelensis* (Bti) bacterial larvicide spray applications have become important aspects of integrated vector control programs for mosquito control^45^, suggesting that development of spray RNAi yeast formulations could be successfully pursued.

As the potential efficacy of RNAi ATSBs continues to be evaluated, it will also be critical to conduct engagement activities with relevant stakeholders. Such activities, which were performed in conjunction with mosquito yeast RNAi field trials^46^, will promote trust and cooperation between farmers and scientists, build a community united to combat SWD, allow stakeholders to provide input into product development, and increase access to the new technology by identifying farms to participate in future scaled product testing^47^. Through early and ongoing stakeholder engagement, a highly effective, stakeholder-accepted new SWD control product could be produced. It is anticipated that yeast ATSB pesticides, which are projected to be cost-competitive, can be readily produced at scale, and are environmentally-friendly^19,40^, will help promote sustainable SWD control.

## 5. Conclusions

The findings of this study demonstrate that a species-specific RNAi yeast insecticide can serve as a highly toxic component of an ATSB that effectively kills *D. suzukii,* an invasive fruit and berry pest. The yeast ATSB, which silences the *Sh* gene resulting in severe neural and behavioral phenotypes, can be prepared using soda delivered in an inexpensive easily constructed soda bottle feeder which effectively killed SWD under simulated field conditions. RNAi yeast insecticides may represent a new class of effective, yet biorational insecticides that can be used in integrated pest management programs for control of this destructive insect pest.

## Acknowledgements

We thank Juliana Wilson and Juang Huang for providing the *D. suzukii* strain and for useful discussions. We thank members of the lab for comments. This work was funded by USDA NIFA Award 2024-70006-43410.

---

**Supp Fig 1.** Steps demonstrating construction of the YES feeder prepared for delivery yeast-soda ATSBs to *D. suzukii*.

