## Supplementary figures and images for "Development of an eco-friendly RNAi yeast attractive targeted sugar bait that silences the *Shaker* gene in spotted-wing drosophila, *Drosophila suzukii*"

### Supp. Fig. 1

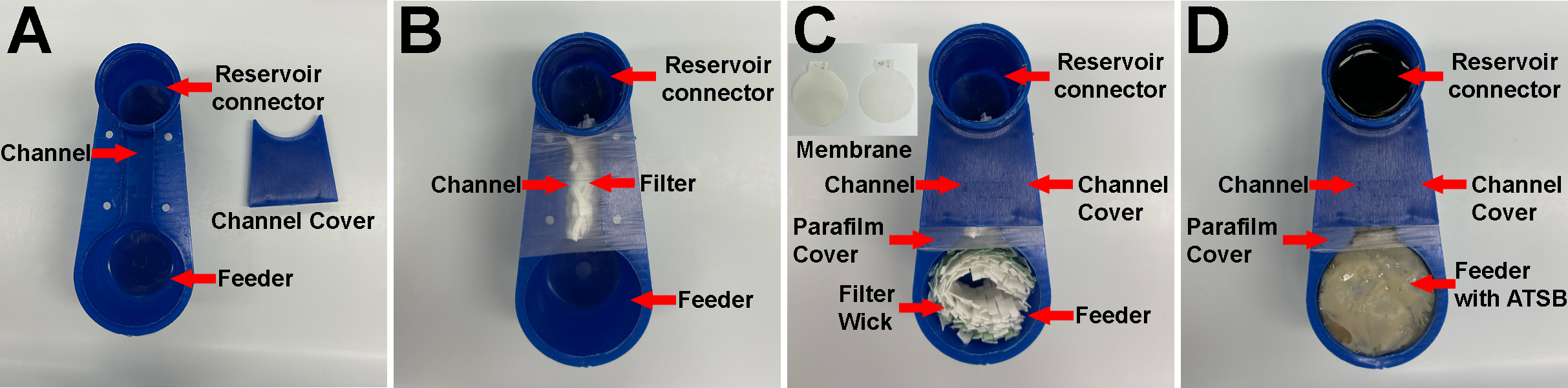
